# Physicochemical characteristics and high sensory acceptability in cappuccinos made with jackfruit seeds replacing cocoa powder

**DOI:** 10.1101/317313

**Authors:** Fernanda Papa Spada, Paula Porrelli Moreira da Silva, Gabriela Fernanda Mandro, GregÓrio Borghese Margiotta, Marta Helena Fillet Spoto, Solange Guidolin Canniatti-Brazaca

**Author notes:** E-mails and Orcid, 0000-0001-6885-1870, 0000-0001-9219-6343, 0000-0001-5397-7112 (Fernanda Papa Spada). Av. Laudelina Cotrin de Castro, 271. BQ Água Branca. Piracicaba – São Paulo – Brazil. CEP 13425-110.

## Abstract

Jackfruit seeds are an under-utilized waste product in many tropical countries. In this work, we demonstrate the potential of roasted jackfruit seeds to substitute for cocoa powder in cappuccino formulations. Two different flours were produced from a hard variety jackfruit by drying or fermenting the seeds prior to roasting. Next, seven formulations were prepared with 50%, 75%, and 100% substitution of cocoa powder with jackfruit seed flours. The acceptance of cappuccinos by consumers (n=126) and quantitative descriptive analysis (QDA^®^) were used to describe the preparations. Physicochemical proprieties were also evaluated. When 50% and 75% cocoa powder was replaced with dry jackfruit seed flour, there was no change in sensory acceptability or technological proprieties; however, it is possible identify advantages to using dry jackfruit seed flour, including moisture reduction and high wettability, solubility and sensory acceptation of the chocolate aroma. The principal component analysis of QDA^®^ explained 90% variances; cluster analysis enabled the definition of four groups for six cappuccino preparations. In fact, dry jackfruit seed flour is an innovative cocoa powder substitute; it could be used in food preparations, consequently utilizing this tropical fruit waste by incorporating it as an ingredient in a common product of the human diet.

## 1. Introduction

Jackfruit (*Artocarpus heterophyllus Lam.*) is a syncarp native to India, that is present in tropical regions and composed of stuffs pulp and seeds (1). Ripe jackfruits are large, with weights ranging from 2 to 36 kg; seeds represent 18%–25% the fruit weight (2). Generally, jackfruit is eaten raw and processed (canned juice and leather), and seeds are eaten after boiling, steaming and roasting (2,3). However, jackfruit is still underused due to seasonality, difficulty in logistics and conservation, and low consummation due to a high sensory intensity of taste and aroma in addition to an association of jackfruit with poor communities. Thus, jackfruit is seldom added to other products.

Jackfruit seeds are a source of fiber, potassium, calcium and sodium (3,4). In recent years, jackfruit seeds gained the attention of researchers as an alternative source of starch and protein that can be industrially exploited (2,3,5). In addition, roasting jackfruit seeds (after drying and/or fermentation processes) produced changes in the aroma sensory profile that resulted in an agreeable chocolate aroma. The main final volatile composition in jackfruit seeds included pyrazines, Strecker aldehydes, alcohols, esters and furanes (6).

Jackfruit seed flour has the added advantage of less calories than cocoa due to the lower lipid compositions of jackfruit seeds (0.7%–2.2%) compared to Forastero cocoa beans (53%–39%) (7,8). Recently, the price of cocoa has climbed, so there is an incentive in the food industry to find a cocoa substitute (9). In addition, the estimates of cocoa beans demand by 2020 is large; however, production is not expected to grow significantly in the next 10 years (10).

Generally, studies with cocoa substitutes formulate chocolate with milk (11–13). Cappuccino is a potential product that uses milk as an ingredient because consumers expect a chocolate aroma without strong chocolate taste. This study characterized and developed a food application for jackfruit seed flour and consequentially, utilizes this tropical fruit waste by incorporating it as an ingredient in a common product of the human diet.

## 2. Materials and Methods

### 2.1. Jackfruit

Jackfruits, hard pulp variety, was manually collected in the countryside of São Paulo, Brazil, and fruits of similar size (5 ± 1 kg) and maturity. Jackfruits were cleaned manually in running water, and the seeds were removed. Cocoa powder was provided by Cargill^®^. The other ingredients were bought at a local market.

### 2.2. Jackfruit seed flour

According to (6) two different treatments were executed. For dry jackfruit seed flour: seeds were dried in an oven at 60°C with air circulation, for 48 h. Dry seeds were roasted in a rotary electric oven for 47 min at 171°C. For fermented jackfruit seed flour, seeds were fermented with pulp and banana leaves, the fermented seeds were dried and roasted at 154°C for 35 min.

### 2.3. Cappuccino formulations

The formulations were elaborated based on American patent US5721003A (14), ingredients of Melitta^®^ cappuccino, Nescafé^®^ cappuccino capsule, the Brazilian food legislation and ingredients recommendation (15). They produced seven kinds of cappuccinos: one control with 15% to cocoa powder, three products that replaced cocoa powder with 3.75%, 7.5% and 11.25% for dry jackfruit seed flour and three others uses fermented flour in the same perceptual (Fig. 1A).

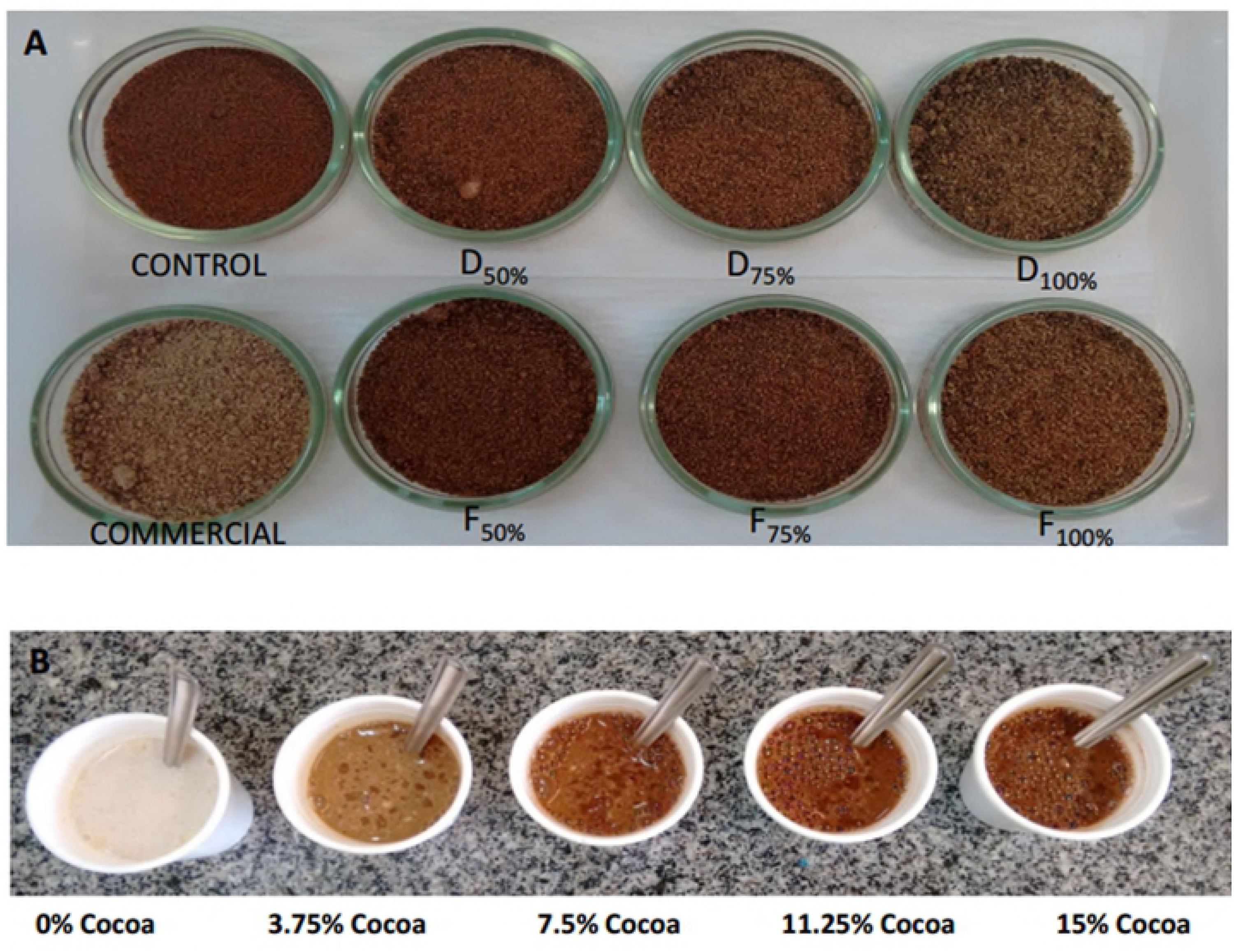
Figure 1A. Cappuccino formulations. Control: produced with 15% cocoa powder; dry jackfruit seed flours (D); and fermented jackfruit seed flours (F). Proportions with 50%, 75% and 100% substitution. Commercial: Melitta^®^ powder for traditional cappuccino formulation. Figure 1B. Cappuccino preparations for QDA^®^ reference scale extremes. Cappuccino with 0.00%; 3.75%; 7.50%; 11.25% and 15.00% cocoa powder.

### 2.4. Physicochemical analysis

Water activity was measured from the temperature of the dew point (Aqualab^®^), and moisture was determined by a standard gravimetric method using infrared light (Bel Engineering Modelo B-TOP-Ray). The pH was determined using two grams of the cappuccino formulation added to 20 mL distilled water. The wettability was measured using the immersion method based on the work of (16). The time between placing a powder sample (2.5 g) of given height (5 cm) on a liquid surface (80°C) and achieving complete wetting was determined. The measurement enabled free sinking of particles, so that unsteady and steady state wetting occurred (16). For calculations, the equation one (Eqn 1) was applied. The apparent density was measured, based on the work of (17), in a 100 mL graduated cylinder, by addition of the required weight of the cappuccino formulation to generate 30 mL.

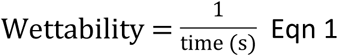

Solubility was determined based on the method used by (5) by weighing 0.1 g cappuccino formulations into weighed centrifuge tubes and adding 10 mL of distilled water. The suspension was stirred and placed in a water bath for 20 min at 80°C, and then the tubes were centrifuged (NT 825) for at 25°C for 15 min at 4420 rpm. An aliquot (5 mL) was transferred from the supernatant to a Petri dish and placed on the stove at 105°C for 24 h to determine the weight of the solid. The solubility was determined by Eqn 2:

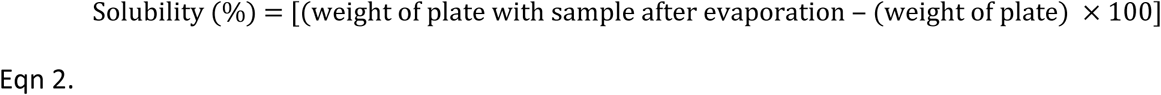

#### 2.4.1. Instrument color analysis

Color was measured using a Minolta^®^ colorimeter, with illuminant C, previously calibrated with a white surface (Y = 93.7, x = 0.3135 and y = 0.3195) based on the CIE-lab L*, a* and b* scale.

### 2.5. Sensory analysis

#### 2.5.1. Sample preparation

The cappuccino formulations were portioned into 10 g samples. During the sensory tests, 50 mL water at 60°C was added to cappuccino formulations in a polystyrene thermal cup. Every sample was prepared immediately after the panelist arrived.

#### 2.5.2. Consumer study

The acceptance test was carried out on a laboratory scale (18,19) with one session using 126 consuming assessors (55% female; 18–42 years old; nonsmoker), selected because they liked and consumed coffee. The consuming assessors evaluated the appearance, aroma, taste and overall impression using a nine points hedonic scale (1=disliked extremely; 9=liked extremely). Samples were randomly evaluated; the consumers were considered repetitions in an incomplete block reply 18 times (T = 7, k = 4, r = 4, B = 7, L = 2, E = 0.88). Where T is number of samples; k is number of samples in each ranking test; r is number of times each sample was shown in each block; B is number of panelists in each block; L is number of times the samples were shown together; and E is dependability of the analysis (20).

#### 2.5.3. QDA^®^

Sensory evaluations were approved by the Ethics Committee of Human Research (COET/077/131). Conventional profiling using QDA^®^ was applied according to (18,19,21). In the first stage, 20 nonsmoker volunteers were recruited and a pre-selection was performed to evaluate their ability to discriminate tastes and odors through basic taste tests. For the second stage, 12 panelists were selected, all females, 18–35 years old, to define the descriptive terminology for the sensory attributes of cappuccino with dry or fermented jackfruit seeds during six training sessions.

Each attribute was provided, together with definitions and physical references using formulated cappuccinos, similar to commercial and cappuccinos added to jackfruit seed flours (Fig. 1B).

The generation of a unique list of attributes was achieved by consensus, and the discrepant terms were eliminated. The final attributes were chocolate (choaro), cappuccino (caparo), coffee (cofaro), cinnamon (cinaro), and fermented (feraro) as attributes for aroma; chocolate (chotas), cappuccino (captas), and fermented (fertas) as attributes for taste; brown (broapp) as the attribute for appearance; gritty (gritex) as the attribute for texture; and overall impression (oveimp). Thus, the reference material was established and the intensity scores were determined for each attribute (Table 1), which were used in the sensory analysis stage.

**Table 1.**
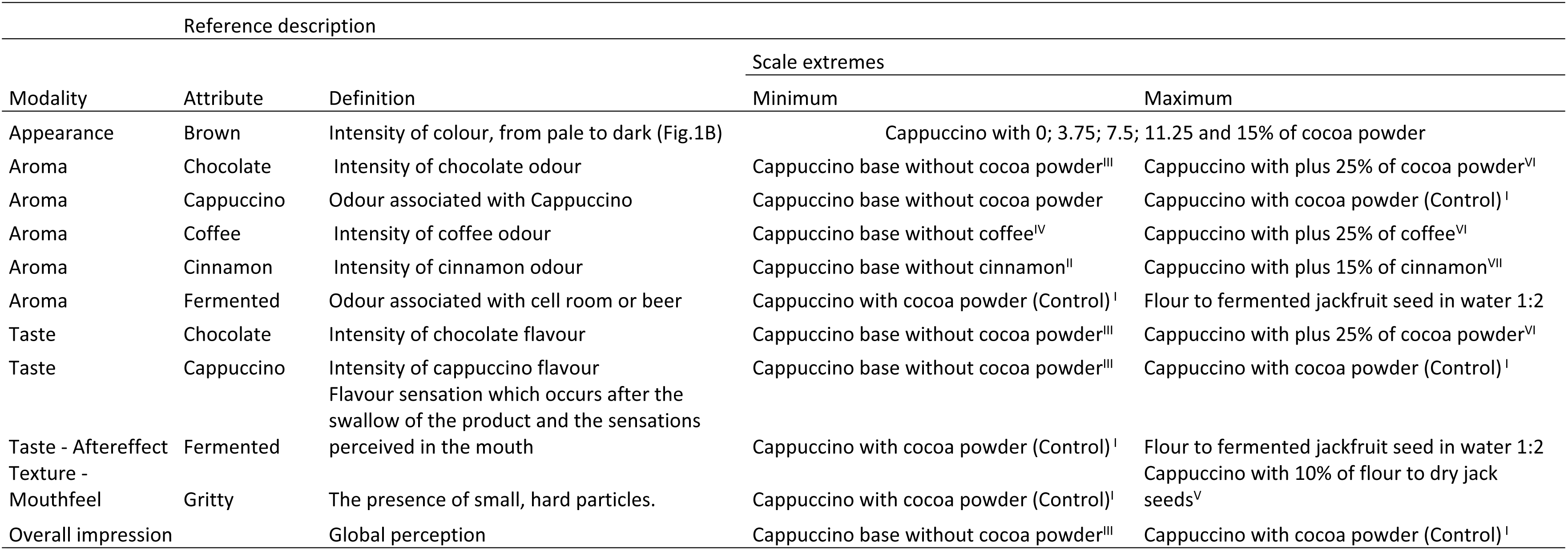
QDA^®^ attributes, definitions and descriptions for standard. Dry seeds flour (D); Fermented seeds flour (F). Proportions with 50, 75 and 100% of substitution.

The sensorial evaluation of the samples was performed in three sessions for cappuccinos with dry jackfruit seeds and another three sessions for cappuccinos with fermented seeds. The tasters were instructed to describe the sensations perceived regarding all final attributes of the samples using a nine-point intensity scale ranging from less intense to more intense for attributes.

### 2.6. Statistical analysis

Physicochemical analyses were available in triplicate, and analysis of variance (ANOVA) was carried out to analyze the results. The comparisons of treatments were performed with Tukey’s test (p≤0.05). The acceptance test was determined using Compusense Five^®^ by the Tukey’s test (p≤0.05). The QDA^®^ results were submitted to multivariate analysis using the correlation analysis (CORR), principal component analysis with biplot graph (PCA) and cluster analysis (CA). In the CA, the cutoff was the average method with the Euclidean distance as the similarity coefficient with a cutoff at |0.70|.

## 3. Results

### 3.1. Physicochemical analysis

The density was similar (p≤0.05) for all available treatments (Table 2). The addition of more than 50% fermented jackfruit seed flour reduced the pH compared to the cappuccino formulated with cocoa powder. Moisture and aW were lowest in cappuccino made with jackfruit seeds. Wettability and solubility were higher in cappuccino with jackfruit seed flour compared to the control (Table 2).

**Table 2.** Physicochemical properties, color characterization, of sensory scores of cappuccino preparations. Control: Cappuccino with 15% cocoa powder; proportions with 50%, 75% and 100% substitution. D50: cappuccino with 7.5% dry jackfruit seed flour and 7.5%cocoa powder; D75: cappuccino with 11.25% dry jackfruit seed flour and 3.75% cocoa powder; D100: cappuccino with 15% dry jackfruit seed flour; F50: cappuccino with 7.5% fermented jackfruit seed flour and 7.5% cocoa powder; F75: cappuccino with 11.25% fermented jackfruit seed flour and 3.75% cocoa powder; and F100: cappuccino with 15% fermented jackfruit seed flour. Different letters in the same column differ significantly (p≤ 0.05) by Tukey’s test.

|  | Control | D50 | D75 | D100 | F50 | F75 | F100 | Commercial |
| --- | --- | --- | --- | --- | --- | --- | --- | --- |
| *Physicochemical properties* |  |  |  |  |  |  |  |  |
| aW | 0.39 ± 0.004<sup>a</sup> | 0.38 ± 0.001<sup>b</sup> | 0.36 ± 0.003<sup>c</sup> | 0.33 ± 0.007<sup>f</sup> | 0.37 ± 0.001<sup>c</sup> | 0.38 ± 0.001<sup>b</sup> | 0.35 ± 0.001<sup>d</sup> | - |
| Moisture | 3.35 ± 0.16<sup>a</sup> | 2.84 ± 0.04<sup>ab</sup> | 3.12 ± 0.08<sup>ab</sup> | 3.09 ± 0.02<sup>ab</sup> | 2.83 ± 0.04<sup>ab</sup> | 2.75 ± 0.05<sup>ab</sup> | 2.77 ± 0.04<sup>ab</sup> | - |
| pH | 6.76 ± 0.005<sup>ab</sup> | 6.75 ± 0.002<sup>b</sup> | 6.71 ± 0.002<sup>b</sup> | 6.82 ± 0.002<sup>a</sup> | 6.73 ± 0.003<sup>b</sup> | 6.55 ± 0.002<sup>c</sup> | 6.56 ± 0.003<sup>c</sup> | - |
| Wettability | 0.07 ± 0.06<sup>c</sup> | 0.28 ± 0.10<sup>b</sup> | 0.20 ± 0.03<sup>bc</sup> | 0.29 ± 0.02<sup>b</sup> | 0.32 ± 0.10<sup>b</sup> | 0.24 ± 0.04<sup>bc</sup> | 0.39 ± 0.04<sup>ab</sup> | - |
| Apparently density | 17.93 ± 0.02<sup>a</sup> | 19.01 ± 0.04<sup>a</sup> | 18.64 ± 0.04<sup>a</sup> | 18.92 ± 0.02<sup>a</sup> | 18.18 ± 0.01<sup>a</sup> | 18.74 ± 0.02<sup>a</sup> | 18.46 ± 0.00<sup>a</sup> | - |
| Solubility | 3.08 ± 0.243<sup>de</sup> | 3.77 ± 0.081<sup>ab</sup> | 3.71 ± 0.015<sup>bc</sup> | 2.97 ± 0.210<sup>e</sup> | 3.53 ± 0.312<sup>bcde</sup> | 3.58 ± 0.225<sup>bcd</sup> | 3.19 ± 0.031<sup>cde</sup> | - |
| *Color* |  |  |  |  |  |  |  |  |
| Lightness (L*) | 40.67 ± 0.19<sup>e</sup> | 44.07 ± 0.37<sup>d</sup> | 47.40 ± 0.55<sup>c</sup> | 52.48 ± 0.62<sup>b</sup> | 44.37 ± 0.33<sup>d</sup> | 44.99 ± 0.55<sup>cd</sup> | 51.30 ± 2.68<sup>b</sup> | 61.09 ± 0.41<sup>a</sup> |
| Redness (a*) | 11.02 ± 0.17<sup>a</sup> | 10.35 ± 0.06<sup>b</sup> | 9.72 ± 0.17<sup>c</sup> | 8.17 ± 0.12<sup>d</sup> | 10.53 ± 0.16<sup>b</sup> | 9.41 ± 0.12<sup>c</sup> | 8.14 ± 0.42<sup>d</sup> | 7.99 ± 0.12<sup>d</sup> |
| Yellowness (b*) | 11.66 ± 0.17<sup>ab</sup> | 11.71 ± 0.1<sup>ab</sup> | 11.73 ± 0.11<sup>ab</sup> | 11.23 ± 0.07<sup>b</sup> | 11.89 ± 0.18<sup>a</sup> | 11.53 ± 0.08<sup>ab</sup> | 11.63 ± 0.72<sup>ab</sup> | 7.33 ± 0.11<sup>c</sup> |
| Chroma (C) | 16.05 ± 0.01<sup>a</sup> | 15.63 ± 0.01<sup>abc</sup> | 15.23 ± 0.01<sup>bc</sup> | 13.89 ± 0.01<sup>e</sup> | 15.88 ± 0.02<sup>ab</sup> | 14.89 ± 0.01<sup>cd</sup> | 14.19 ± 0.06<sup>de</sup> | 10.84 ± 0.01<sup>f</sup> |
| Hue (H°) | 46.62±0.00<sup>e</sup> | 48.51±0.01<sup>d</sup> | 50.35±0.01<sup>c</sup> | 53.96±0.01<sup>b</sup> | 48.49±0.00<sup>d</sup> | 50.78±0.01<sup>c</sup> | 55.01±0.01<sup>a</sup> | 42.56±0.01<sup>f</sup> |
| *Sensory acceptance* |  |  |  |  |  |  |  |  |
| Appearance | 7.29<sup>a</sup> | 7.05<sup>a</sup> | 7.06<sup>a</sup> | 6.99<sup>a</sup> | 7.10<sup>a</sup> | 7.19<sup>a</sup> | 7.06<sup>a</sup> | - |
| Aroma | 6.74<sup>bc</sup> | 7.55<sup>a</sup> | 7.04<sup>ab</sup> | 6.46<sup>bc</sup> | 6.06<sup>cd</sup> | 5.56<sup>d</sup> | 5.62<sup>d</sup> | - |
| Taste | 6.60<sup>a</sup> | 6.54<sup>a</sup> | 6.31<sup>ab</sup> | 5.95<sup>abc</sup> | 5.65<sup>bc</sup> | 4.51<sup>d</sup> | 5.12<sup>cd</sup> | - |
| Overall impression | 6.83<sup>a</sup> | 6.88<sup>a</sup> | 6.71<sup>a</sup> | 6.30<sup>ab</sup> | 6.02<sup>bc</sup> | 5.30<sup>d</sup> | 5.54<sup>cd</sup> | - |

### 3.2. Instrumental color and consumer study

Color results demonstrated the 50% cocoa substitution had similar chroma values compared to the cappuccino control (p≤0.05) (Table 2). Independent of the kind of jackfruit seed flour, 75% produced formulations with the same yellowness (b*) as the control, but the clearest were observed with Hue results (Table 2). The commercial standard was the clearest (low L*, chrome and Hue) (Table 2 and Fig. 1A).

Other evidence of similar color was related by consumers, because they not only found appearance modifications between preparations (p≤0.05), but the values indicated greater acceptability, with scores higher than seven (Table 2). In fact, cappuccino color did not change the sensory acceptation.

The dry jackfruit seed flour utilization with 50% and 75% substitution improved the aroma and cappuccino acceptability. However, fermented flour used for 75% and 100% cocoa powder substitution reduced the aroma acceptability (Table 2). The taste was still similar to the control using dry jackfruit seed flour (p≤0.05), but fermented flour reduced the acceptance of taste (Table 2). The overall impression was still equal to the control when dry jackfruit seed flour was used (p≤0.05).

### 3.3. Quantitative descriptive analysis (QDA^®^)

#### 3.3.1 Correlation analysis (CORR)

In the CORR analysis, based on the method of (22), values above |0.70| were consider accented, and 22 correlations were thus selected. There are seven above 0.90, which were regarded as very strong, and there are 13 others above|0.70|. All variables had at least one correlation; for caparo, strong correlations were observed with captas, broapp, cinaro and oveimp.

#### 3.3.2. Principal component analysis (PCA) and cluster analysis (CA)

The PC1 explained 62% of the statistical variance and was positively correlated (right side) with the variables caparo, captas, chotas, cinaro and cofaro and negatively correlated (left side) with feraro and fertas. The PC2 explained 28% of the statistical variance and was positively correlated (top) with the variables broapp and feraro and negatively (bottom) correlated with gritex (Fig. 2).

**Fig. 2.**
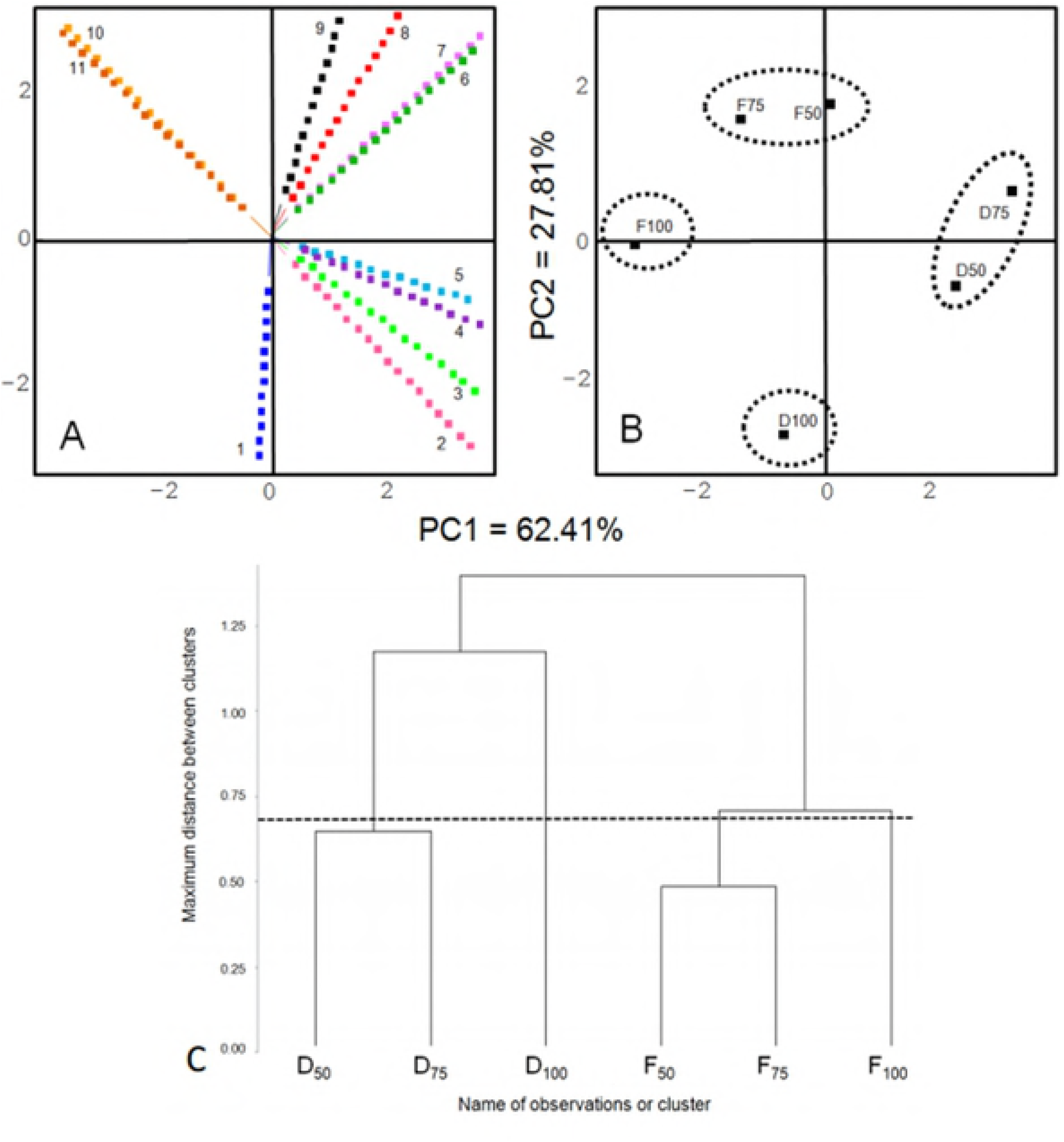
Projection of variables (A), group observations (B) and Dendrogram from the QDA^®^ results of cappuccino formulations (C). Principal component analysis using the sensory attributes of cappuccinos formulated with jackfruit seed flours. PC1 and PC2: principal components 1 and 2. 1- gritex: gritty texture; 2- oveimp: overall impression; 3-cofaro: coffee aroma; 4- cinaro: cinnamon aroma; 5- chotas: chocolate taste; 6- captas: cappuccino taste; 7- caparo: cappuccino aroma; 8- broapp: brown appearance; 9- choaro: chocolate aroma; 10 -feraro: fermented aroma; and 11- fertas: fermented taste. Dry seed flours (D); fermented seed flours (F). D50: cappuccino with 7.5% dry jackfruit seed flour and 7.5% cocoa powder; D75: cappuccino with 11.25% dry jackfruit seed flour and 3.75% cocoa powder; D100: cappuccino with 15% dry jackfruit flour; F50: cappuccino with 7.5% fermented jackfruit seed flour and 7.5% cocoa powder; F75: cappuccino with 11.25% fermented jackfruit seed flour and 3.75% cocoa powder; and F100: cappuccino with 15% fermented jackfruit seed flour.

In CA, observations were separated into four groups (D100; D50 and D75; F100; and F50 and F75) by the cutoff held at |0.70|, shown by the dotted line. The same cutoff was used to separate the groups (dotted circles) shown in the projection of observations (Fig. 2B). Generally, treatments D100 and F100, received the lowest scores compared to other groups (D50-D75 and F50-F75). In F75 group, low coffee, cappuccino and cinnamon aromas and chocolate and cappuccino tastes were noted. The appearance was light brown, and the fermented tastes and aromas were more evident. The group formed by D50 and D75 had high caparo, captas, choaro, chotas, cinaro and cofaro.

## 4. Discussion

The use of jackfruit seed with a replacing of cocoa powder in cappuccino formulations is possible, dry seeds have more potential because it not has taste and aroma fermented. The ideal level of cocoa substitution in cappuccino formulations are nearest 50 and 75%, it is possible find some vantages using jackfruit seed flour, as a moisture reduce, high wettability, solubility, sensory acceptation by chocolate aroma and other similarities, such as color, density, pH, aW and appearance. Fermented attributes under-characterized the cappuccino preparations; PCA explaining 90% variance; using CA was possible define four groups to seven cappuccino preparations.

### 4.1. Physicochemical analysis

The pH would affect the intensity of the sour taste, which is linearly related to the summation of the molar concentrations of organic acid species that contain at least one protonated carboxyl group plus the concentration of free hydrogen ions (23). In addition, salt of the organic acid in the food further lowers the ionization by common effect (24). In cocoa nibs, the alkalization process raises the pH. This process provides cocoa nibs at pH 6.0 with chocolate flavor or nibs at pH 7.2 to 8.1 that is typical of dark chocolate with the sour, bitter, fruity and moldy characteristics. This may influence the acceptability reductions in cappuccinos with fermented jackfruit seed flour.

Formulated cappuccinos have aW similar to that found by (13) when they developed milk beverages with some varieties of carob powder. The values of aW were between 0.29 and 0.41. However, (25) identified 20% higher aW using carob powder, and moisture was three times larger in comparison to cappuccinos: 9.6% and 9.0% for carob and cupuassu, respectively. These are technological advantages because they reduce potential microorganism proliferation and change compaction and mechanical proprieties (26).

The low wettability in the cappuccino control was influenced by high cocoa concentration (27), and this value was compatible to (11). Cappuccino formulations had higher wettability, which predisposed for higher levels of solubility, probably because cocoa beans have ten times more lipids than jackfruit seeds. On effect, cappuccinos with jackfruit seeds are similar or better than other natural substitutes currently in use (carob and cupuassu).

### 4.2. Instrumental color and consumer studies

The dark brown color in jackfruit seeds was produced by the Maillard reaction, also typical in cocoa nibs due to ideal roasting conditions, reducing sugars and amino groups (7,8,28). In this study, it was independent of the fermentation process. The color in cappuccinos with jackfruit seeds did not change the sensory acceptance; thus, cappuccino formulations with jackfruit seeds flour had compatible or better color in comparison to control and commercial cappuccinos, which is also another indication of the high quality of this natural cocoa substitute sample.

These results demonstrated the innovate potential of dry jackfruit seeds as a cocoa powder replacer. Carob and cupuassu are established substitutes for cocoa powder; however, when it was used in milk beverages, the sensory acceptation was reduced (11,12). Before this study, the low acceptability of cocoa substitutes was justified due to low lipid concentrations, but jackfruit seeds also have low lipids and do not reduce the consumer acceptance when substituted for 50% or 75% cocoa powder. Most likely, the pH reduction in cappuccino using fermented flours (75% and 100% substitution) improved sour and moldy tastes; this changed the acceptability significantly (p≤0.05) for taste, aroma and overall impression.

### 4.3. Quantitative descriptive analysis (QDA^®^)

#### 4.3.1. Correlation analysis (CORR)

In a new cappuccino preparation, the aroma and taste characteristic were interdependent with brown color and cinnamon aroma as a positive impact on overall impressions. Briefly, the aromas of cappuccino, coffee, cinnamon, and the taste of cappuccino were highly correlated and expected for a good overall impression of cappuccinos. The fermented aroma and taste were very interdependent and not characteristic in these preparations. This explained the few overall impressions of cappuccinos made with fermented jackfruit seed flour.

#### 4.3.2. Principal component analysis (PCA) and cluster analysis (CA)

The choaro was higher in F100, which is possible because the fermentation process improves volatile compounds, such as pyrazines and esters (6), and because dark color is generally associated with high chocolate concentration, meaning that broapp was great in F100. However, the high values for feraro and fertas in F100 produced an over-taste not characteristic of cappuccinos; thus D100 received more caparo and chotas in comparison to F100, even when D100 was less soluble.

The other group for CA corresponding to F50 and F75 was characterized as little gritex, caparo, cofaro and chotas; medium cinaro and higher capta, browapp, choaro, fertas and feraro. Thus, even the high chocolate aroma of the characteristic cappuccino aroma was reduced because other fermented flavors were found; these flavors were not expected in cappuccino preparations. The recognized food was related to the construction of different sensory systems to name foods or first to cause our survival (29). To minimize the fermented flavor, appropriate amounts of flavors could be produced depending on fermentation conditions. For example, the mode of fermentation, natural or inoculate, which is usual for cocoa beans (8,30). In fact, better standardization of the chocolate fermentation process could be generate a chocolate with different sensory characteristics (31). Thus, future studies will know the flavor characteristics of jackfruit seeds using inoculated cultures of microorganisms.

## 5. Conclusions

Dry jackfruit seed flour can be incorporated as an ingredient in cappuccino formulations; 50% and 75% substitution of cocoa powder by dry jackfruit seed flour did not change sensory acceptability or characteristics. Fermented attributes were not characteristic of cappuccinos, but they improved the chocolate aroma. The primary characteristics responsible for the character of cappuccinos with dry jackfruit seeds were cappuccino, chocolate, cinnamon and coffee aromas, and cappuccino and chocolate tastes.

